# Wild to domesticates: genomes of edible diploid bananas hold traces of several undefined genepools

**DOI:** 10.1101/2021.01.29.428762

**Authors:** Julie Sardos, Catherine Breton, Xavier Perrier, Ines Van Den Houwe, Janet Paofa, Mathieu Rouard, Nicolas Roux

## Abstract

This study is an unprecedent exploration of the diversity of 226 diploid bananas genotyped with restriction-site-associated DNA sequencing data (RADseq) to clarify the processes that led to the creation of edible diploid AA bananas. This wide set included 72 seedy bananas, mostly *M. acuminata* from different genepools, and 154 edible, i.e. parthenocarpic and sterile, AA accessions obtained from genebanks and recent collecting missions. We highlighted the geographic organisation of the diversity of edible AAs and confirmed the admixed nature of many and further conducted introgressions tests within AAs from South East Asia and New Guinea. Lastly, taking advantage of the presence of an important number of *M. acuminata* ssp. *banksii* (22) and of AA from Papua New Guinea (76) in the set, we investigated the patterns of differentiation between wild and cultivated bananas seemingly belonging to the same genepool. We discovered a few cultivated AAs that may be of pure origins both in South-East Asia and in New Guinea. We also detected two undefined parental genepools in South East Asia for which regions of origin could be Thailand and a region between north Borneo and the Philippines, respectively. Finally, we suggest the existence of a third genepool in New Guinea island that might be a source population for both edible AAs and the local *M. acuminata* ssp. *banksii*.

## Introduction

Bananas, including both dessert and cooking types, are of high importance economically but are also staple for millions of people in developing countries. Often qualified as a giant herb, the genus *Musa* is a monocotyledon that belongs to the order of the Zingiberales and is native to a wide South-East Asia (SEA)/West Oceania region (Janssens et al. 2016). It is divided into two botanical sections, *Musa* (formally *Eumusa* and *Rhodochlamys* sections) and *Callimusa* (formally *Callimusa* and *Australimusa* sections) (Häkkinen 2013). Most edible bananas were domesticated from the *Musa* section, at the notable exception of the rare Fe’i bananas (T genome) that were domesticated from a yet undefined *Callimusa* species. In the section *Musa*, the major and probably earlier ancestor of edible bananas is *M. acuminata* (A genome). *Musa balbisiana* (B genome) and, at a minor extent, other wild species as *M. schizocarpa* (S genome) and undefined Callimusa species (T genome), also contributed to the genetic make-up of current days edible bananas (Heslop-Harrison and Schwarzacher 2007).

In edible bananas, the main traits selected during the wild-to-domesticate transition are parthenocarpy, i.e. the ability to set fruits without the need of prior pollination, and female sterility(Denham et al. 2020). Together, they ensure the production of fleshy fruits that are free of seeds. A side effect of these traits, that are mandatory for consumption, is a vegetative mode of propagation for this crop. Therefore, the diversity of cultivated bananas is fixed over long period of time and the occurrence of new varieties is limited to the clonal fixation of mutations (Noyer et al. 2005; Sardos et al. 2018) and to rare spontaneous sexual events. Nevertheless, in-spite of this mandatory vegetative mode of propagation, millennia of human selection and diversification have led to the current existence of hundreds of edible banana varieties. Among them, the most common and the most widespread are triploids, such as the commercial ‘Cavendish’ (AAA genome group) or the subsistence ‘Plantain’ (AAB genome group). It is less known but edible diploid bananas (AA, AB or rarely AS and AT genome groups) are also cultivated, especially in backyards and subsistence farming systems. They are not very common in Asia but have high cultural importance in certain communities of east Africa and are widely cultivated in Papua New Guinea (Bourke 1976; Nayar 2009; Perrier et al. 2019). In addition, this last country hosts a wide diversity of edible AA bananas, a unique feature in the world (Arnaud and Horry 1997; Sardos et al. 2018; Sardos, Paofa, et al. 2019).

As they gave rise to edible triploids (Raboin et al. 2005; De Langhe et al. 2009), edible diploids could be seen as the earliest banana domesticates, especially those with an AA genomic composition. Under that perspective, a major step towards the complete resolution of banana domestication would go through the comprehension of the transition between *M. acuminata* and edible AAs.

The range of *M. acuminata* spans from India at the west to north Australia at the east. It is divided into several subspecies that are geographically segregated and show unique features, morphologically (Simmonds 1956), genetically (Hippolyte et al. 2012) and at the genome level (Martin, Baurens, et al. 2020). The AA bananas were first hypothesized originating in Malaysia (Simmonds and Shepherd 1955) but molecular studies comparing edible AAs with different *M. acuminata* subspecies later pointed at the Philippines – New Guinea (NG) region as the centre of origin for the crop (Carreel et al. 2002). A domestication model was then further refined using transdisciplinary approaches (De Langhe et al. 2009; Perrier et al. 2009; Perrier et al. 2011). While acknowledging the importance of the NG subspecies *M. acuminata* ssp. *banksii*, this model considers those of the plants selected from a single genepool as cultiwild, i.e. pre-domesticated plants that are not fully parthenocarpic nor sterile. It also emphasizes hybridization between the different subspecies of *M. acuminata* as the main force that drove the rise of fully parthenocarpic, sterile, edible diploid bananas. This model is supported by archaeological evidence of early cultivation, 7000 years ago, of seedy bananas in Papua New Guinea highlands (Denham et al. 2003) and by the recent resolution of complex genomic ancestries in nine present days edible AAs (Martin, Cardi, et al. 2020). However, the genotyping of wide sets of edible diploid banana accessions with microsatellites, DArT and Genotyping-by-sequencing SNPs markers suggested the potential existence of discrete genepools in SEA and NG and questioned the existence of edible diploids AA, fully parthenocarpic and sterile, that would not be products of hybridization, notably in NG (Sardos, Perrier et al. 2016; Sardos, Rouard et al. 2016; Christelova et al. 2017).

In this study, we explored the diversity of 226 samples of diploid bananas genotyped with restrictionsite-associated DNA sequencing data (RADseq) with the aim of clarifying the processes that led to the creation of edible diploid AA bananas.

## Results

### Diversity analysis

The dissimilarity matrix and the NJ Tree obtained on the whole sample set comprising 226 individuals (**Table 1** and **Supplementary Fig. S1**, Supplementary Material online) allowed the identification of 158 unique genotypes including 30 multi-accessions Genotype Clusters. These Genotype Clusters corresponded to duplicates or clonal varieties (**Supplementary Table S1**, Supplementary Material online). The pruned set of 158 accessions kept for further analyses, included 91 edible accessions out of the 154 initially present in the whole dataset. In the NJ tree constructed on the pruned sample, the subspecies of *M. acuminata: banksii, malaccensis, zebrina* and *burmannica/siamea* formed segregated clusters. Three out of the four *M. acuminata* accessions (AMB007, AMB008 and Sup4), collected in Maluku islands (Indonesia, west of NG island), clustered at the margin of ssp. *banksii* while the fourth one AMB004 clustered within ssp. *banksii*. The philippino *M. acuminata* ssp. *errans*, represented by the accession ‘errans’ and ‘UPLB’ initially classified as ssp. *banksii* also clustered at the margin of ssp. *banksii*, along with ‘Borneo’ classified as ssp. *microcarpa*. The subspecies *sumatrana* and *truncata* from Sumatra and the Malay Peninsula respectively, clustered with ssp. *malaccensis* (**Fig. 1C**).

**Table 1:** Origins and classification of the accessions of the sample.

| CLASSIFICATION |  | ORIGIN |  |  |  |  |  |  |  |  |  |  |  |
| --- | --- | --- | --- | --- | --- | --- | --- | --- | --- | --- | --- | --- | --- |
|  | n | PNG | Hawaii | Philippines | Indonesia | Malaysia | Indo /<br>malaysia | Thailand | Vietnam | Myanmar | India | East<br>Africa | Not<br>recorded |
| **AA (CVs)** | **154** | **76** | **0** | **15** | **9** | **12** | **7** | **7** | **3** | **0** | **1** | **22** | **2** |
| **M. acuminata** |  |  |  |  |  |  |  |  |  |  |  |  |  |
| *banksii* | 23 | 20 |  | 1 | 2 |  |  |  |  |  |  |  |  |
| *errans* | 1 |  |  | 1 |  |  |  |  |  |  |  |  |  |
| *zebrina* | 6 |  | 1 |  | 3 |  |  |  |  |  |  |  | 2 |
| *sumatrana* | 1 |  |  |  | 1 |  |  |  |  |  |  |  |  |
| *microcarpa* | 2 |  |  |  | 2 |  |  |  |  |  |  |  |  |
| *malaccensis* | 12 |  |  |  |  | 8 |  | 3 |  |  |  |  | 1 |
| *truncata* | 1 |  |  |  |  | 1 |  |  |  |  |  |  |  |
| *siamea* | 3 |  |  |  |  |  |  | 3 |  |  |  |  |  |
| *burmannica* | 3 |  |  |  |  |  |  |  |  | 3 |  | 4 |  |
| Other seedy | 16 |  |  |  | 6 | 2 |  |  |  |  |  |  | 4 |
| sub-total | **68** | **20** | **1** | **2** | **14** | **11** |  | **6** | **0** | **3** | **0** | **4** | **7** |
| **M. schizocarpa** | **3** | 3 |  |  |  |  |  |  |  |  |  |  |  |
| **M. acuminata x**<br>**M. schizocarpa** | **1** | 1 |  |  |  |  |  |  |  |  |  |  |  |
|  | **226** | **100** | **1** | **17** | **23** | **23** | **7** | **13** | **3** | **3** | **1** | **26** | **9** |

**Fig. 1:**
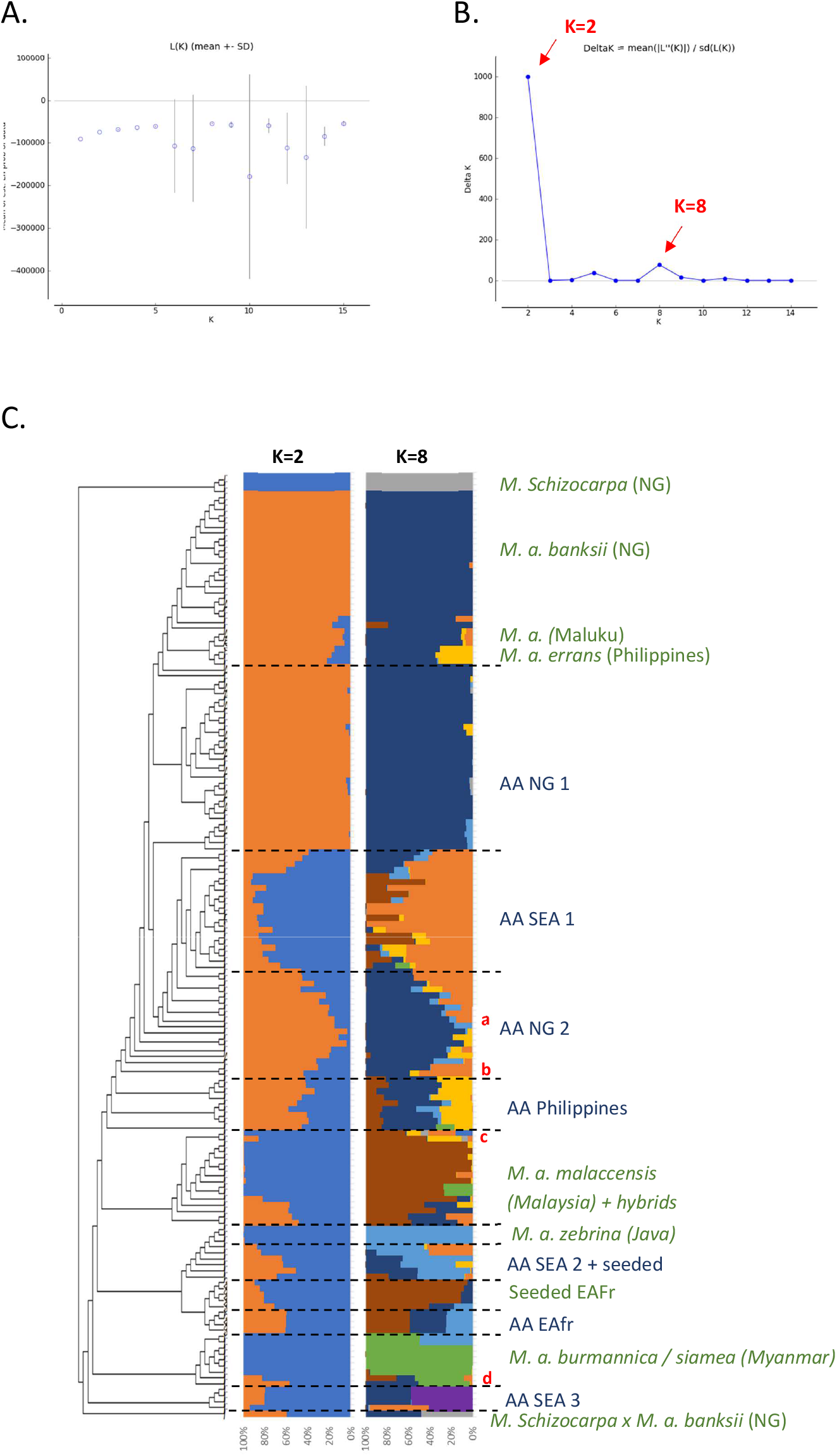
A and B present the results of the Bayesian clustering of the diploid bananas pruned sample (158 accessions) using STRUCTURE v2.3 (Pritchard et al. 2000) evaluated using STRUCTURE HARVESTER (Earl and vonHoldt 2012) and based on the lnP(D)/K and DeltaK, respectively. C presents the global genetic structure of the pruned sample. The cladogram was obtained from the NJ tree constructed by DARwin 6 (Perrier and Jacquemoud-Collet 2006) on the Simple-Matching distance matrix calculated on 66,481 biallelic SNPs and using FigTree v1.4.3 (Rambaut 2006-2016) and the R package *ape* (Paradis and Schliep 2019). Bar plots represent STRUCTURE outputs for K=2 and K=8 as inferred from 1,278 SNPs distributed evenly across the genome, each bar corresponds to a genotype and colours correspond to the detected genetic clusters. EAfr: Africa; SEA: South-East Asia; NG: New-Guinea island; a: ITC0299 ‘Guyod’ from the Philippines, b: ITC0447 ‘Pu-Te Wey’ from Malaysia, c: ITC1701 *M. acuminata* ssp. *sumatrana* from Sumatra and ITC0393 *M. acuminata* ssp. *truncata* from Malaysia, d: ITC1761 ‘Matti’ from India and ITC0610 ‘Tuu Gia’ from Vietnam.

In total, the cultivated AA accessions formed seven clusters (**Fig. 1C**). Among them, a first cluster composed of 31 edible AA accessions from New-Guinea island (denoted ‘AA NG1’), is tightly linked to the *‘banksii’* cluster. Three clusters are not closely related to any subspecies and fall between *‘banksii’* and the wild subspecies from South-East Asia (SEA): ‘AA SEA1’ (15 cultivated accessions from SEA), ‘AA NG2’ (14 cultivated accessions from NG, 1 from the Philippines and 1 from Malaysia) and ‘AA Philippines’ (9 cultivated accessions from the Philippines). The ‘AA SEA2’ (3 cultivated accessions from Indonesia/Malaysia) and ‘AA EAfr’ (5 cultivated accessions from East Africa), were grouped near *‘zebrina’*. We also noted that ‘AA SEA3’ (composed of 3 cultivated accessions classified as Pisang Jari Buaya (PJB) and ‘Pisang Madu’, both from the Indo-Malayan region) clustered apart from any other accessions of our sample. Finally, two accessions, ‘Matti’ and ‘Tuu Gia’ clustered with ‘burmannica/siamea’.

### Population structure

The two best values of k identified by STRUCTURE in the pruned dataset were k=2 and k=8 (**Fig. 1A** and **B**). For k=2, the Bayesian analysis recognized two genepools corresponding roughly to New Guinea (NG) island and South-East Asia (SEA) with a high number of admixed accessions. For k=8, this analysis confirmed discrete genepools for the *M. acuminata* taxa *banksii, malaccensis, zebrina, burmannica/siamea* and *M. schizocarpa*. We noted two accessions from Thailand originally classified as ssp. *malaccensis* that seem to also hold some ssp. *burmannica/siamea* signature in their genome (ITC0668 ‘Pa (Musore) n°2’ with Q _burmannica/siamea_ = 27,1% and ITC1067 ‘THA018’ with Q _burmannica/simaea_ = 27,3%). In addition, the genomic composition inferred for ‘truncata’ and ‘sumatrana’ accessions – belonging to eponymous subspecies - revealed patchworks of different genepools with a *malaccensis* dominance in both. However, their respective genomic profiles are slightly different (**Fig. 1C** and **Fig. 2**).

**Fig. 2:**
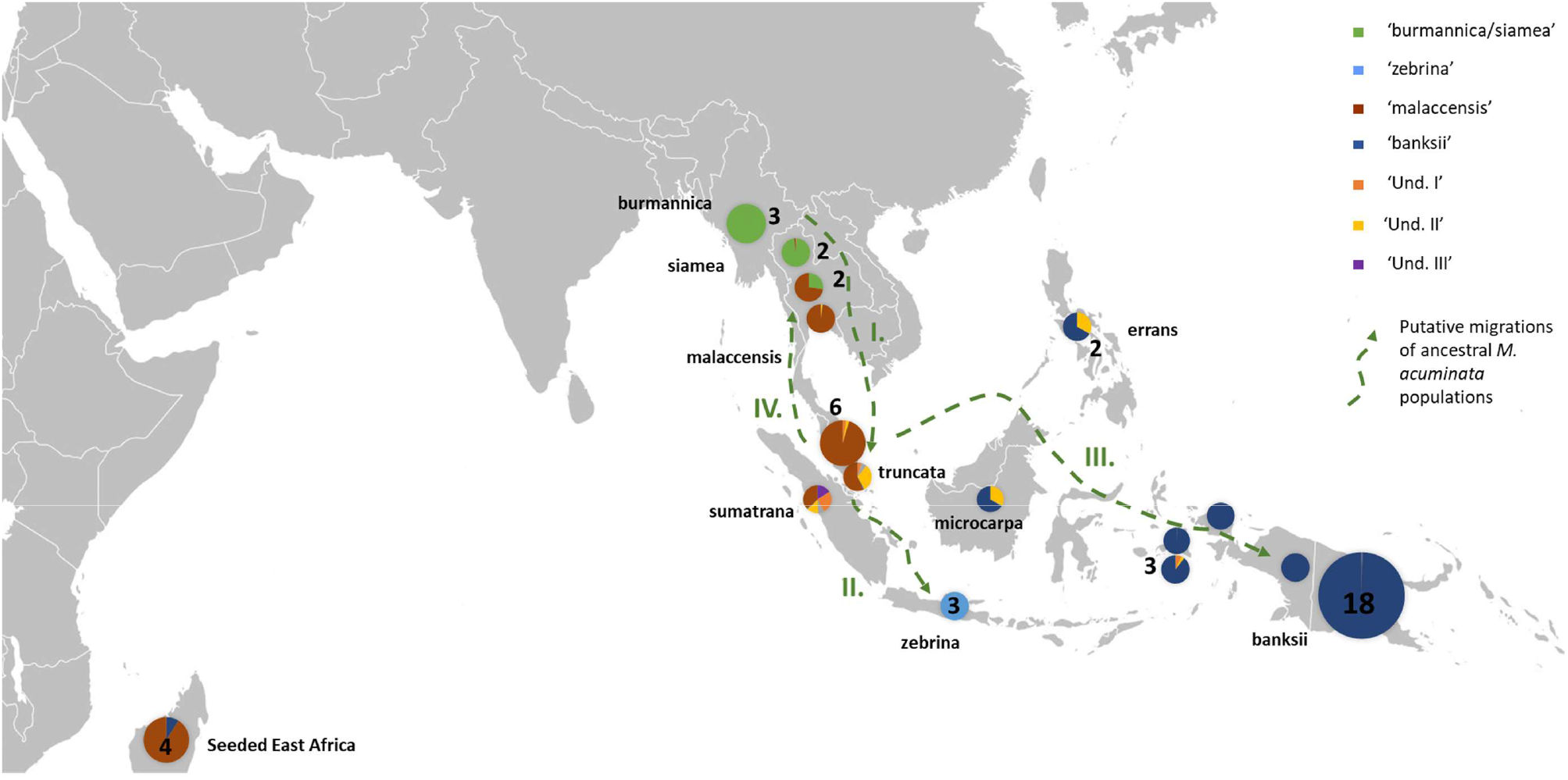
Distribution of the *M. acuminata* accessions of the sample. Pie charts illustrate genomic background as inferred by STRUCTURE v2.3 (Pritchard et al. 2000) for k=8. Numbers indicate the number of accessions sharing similar patterns. Putative migrations of ancestral populations, represented by green dashed arrows, were inferred from Janssens et al. (2016) and Rouard et al. (2018). I. First migration from mainland South-East Asia towards Malayan Peninsula and Sumatra, followed by II. Migration of populations to Java, III. Migrations to Borneo, the Philippines and New-Guinea Island and IV. Secondary migration into mainland South-East Asia.

Several undefined genepools were detected. The first one (denoted ‘Und-I’) (orange colored in **Fig. 1C** and **Fig. 3**) was very common in cultivated accessions. Three cultivars from Thailand and clustering within ‘AA SEA1’, are fully assigned (Q>0.9) to this genepool (‘Thong Dok Mak’, ‘Kluai Lep Mu Nang’ and ‘Sa’). ‘Und-I’ was detected in many cultivated accessions from SEA and in a few accessions of the cluster ‘AA NG2’. It was also inferred as introgressions in some wild specimens spread in different clusters, such as ‘sumatrana’ (Q=0.26) and three hybrids collected in Indonesia NG in the 1960’s and clustering with ‘malaccensis’ (‘Higa’ and ‘Hybrid’) and with ‘banksii’ (‘Waigu’).

**Fig. 3:**
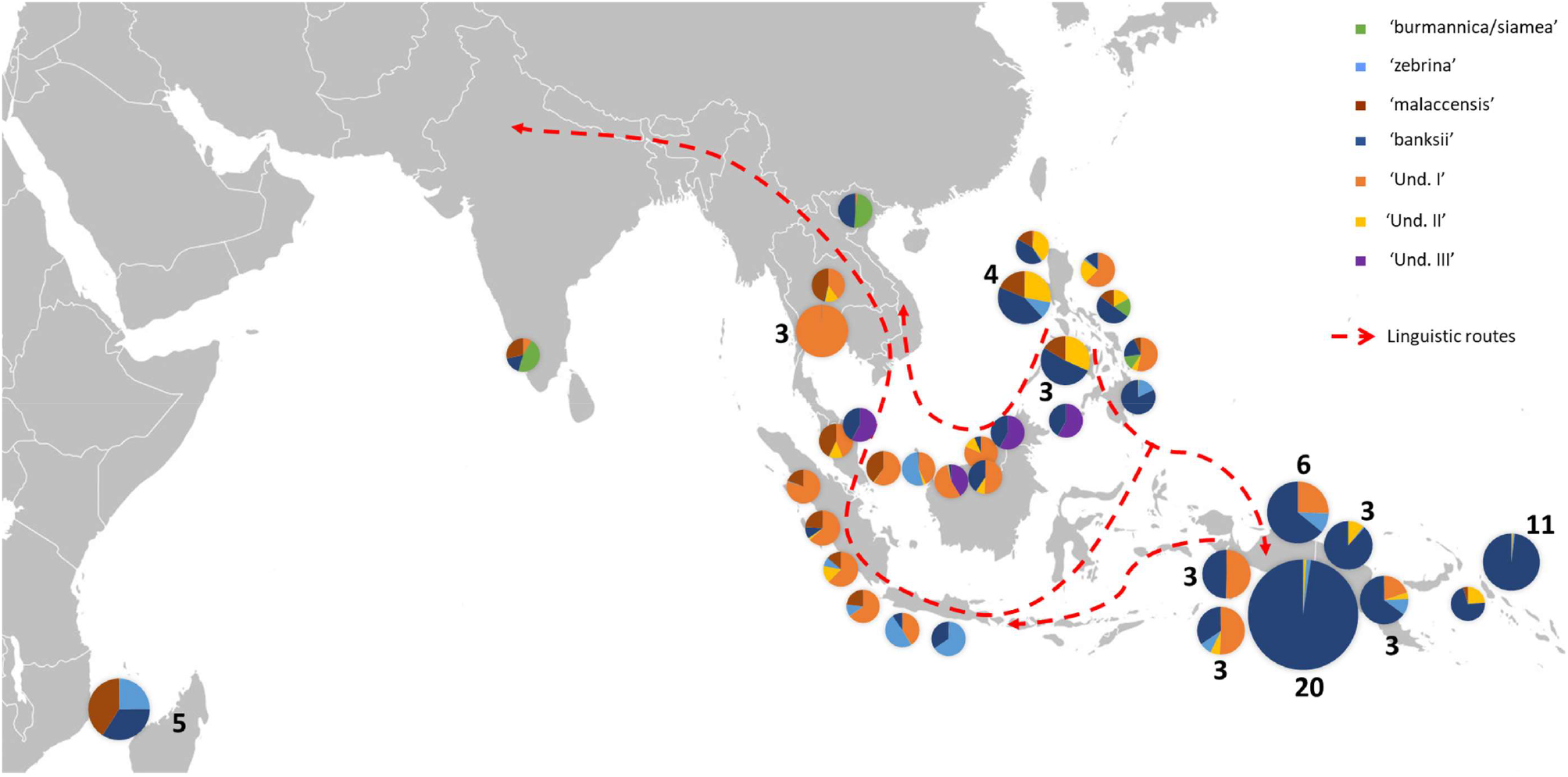
Distribution of cultivated AA accessions of the sample. Pie charts illustrate genomic background as inferred by STRUCTURE v2.3 (Pritchard et al. 2000). Numbers indicate the number of accessions sharing similar patterns. Dashed red lines show historical linguistic routes in the region (Perrier et al. 2011).

The two other undefined genepools were inferred as introgressions only. The genepool ‘Und-II’ was detected in the *errans* subspecies and in the accession ‘Borneo’ who share similar profiles along with a partially common genetic background with ssp. *banksii*. It was also inferred in some of the cultivated accessions including all the ‘AA Philippines’ cluster (yellow color in **Fig. 1C**). The third undefined genepool ‘Und-III’ was identified in the cluster ‘AA SEA3’ (purple color in **Fig. 1C**) in which accessions were collected in the north coast of Borneo and on Sulu, an island located between Borneo and the main Philippines islands (**Fig. 3**). The genepool ‘Und-III’ was also detected as a small introgression in the subspecies *sumatrana*.

The inferred genomic composition of the cultivated AA confirmed the hybrid nature of most of them. *Musa acuminata* ssp. *banksii* from NG is a prominent contributor to AAs, including those in SEA, followed by the Javanese ssp. *zebrina*, ssp. *malaccensis* from the Malayan peninsula and then ssp. *burmannica/siamea*. The latter that ranges from India to north Thailand is the subspecies contributing the less to the edible samples. Interestingly, the genepool ‘Und-II’, detected in the Philippines ssp. *errans*, is also an important contributor, in addition to ‘Und-I’. Finally, we noted in this analysis that some AA accessions in the clusters ‘AA NG1’ and in ‘AA SEA 1’ had a single contributor and did not appear as admixed individuals (**Fig. 1C** and **Fig. 3**).

### Detection of introgressions

#### Of SEA M. acuminata subspecies into AA NG1 (cultivated in NG)

To enable testing the 31 AA accessions of the ‘AA NG1’ cluster for introgression by one or more of the six SE Asian subspecies of *M. acuminata*, we performed 186 tests. Only Patterson’s D tests for which the dominance of the BBAA pattern over ABBA and BABA were considered robust. Following this criteria, ten combinations which all exhibited BABA > BBAA were excluded. Within the remaining 176 tests, Patterson’s D was statistically significantly negative (Z score < −2) for 70 combinations, showing significant excess of BABA sites over ABBA sites and indicating a highest proximity between ssp. *banksii* and the SEA subspecies tested. At the contrary, for 11 combinations, the D scores obtained were significantly positive (Z score > 2), revealing a significant bias towards ABBA pattern compared with BABA and reflecting possible introgression of given SEA subspecies within the AA accessions tested. For 95 combinations tested, D was not significantly departing from 0 (−2 < z-score < 2), therefore not showing significant differences between the counts of ABBA and BABA sites. Among those tests, nine accessions did not showcase any significant differences in the number of ABBA and BABA sites for any of the six SE Asian subspecies tested, suggesting that they may be unadmixed cultivated accessions (**Supplementary Table S2,** Supplementary Material online).

#### Of banksii into AA from SEA

We also performed Patterson’s D tests on 33 AA accessions originating in SEA to check for their introgression by *M. acuminata* ssp. *banksii*. However, for most of the tests, the count of the different patterns showed topology discordance compared to assumption. That is to say that for eight accessions, both ABBA and BABA counts were dominant over BBAA, that for 18 accessions ABBA counts were dominant over BBAA and for one accession, namely ‘Malaysian Blood’, BABA was dominant over BBAA. For all the six tests for which no topology discordance was identified, statistically significant bias towards ABBA pattern were identified when compared to BABA, suggesting introgression of ssp. *banksii* in the accessions tested (**Supplementary Table S3,** Supplementary Material online).

### Pattern of differentiation between *M. acuminata ssp*. banksii and the cluster ‘AA NG1’

Heterozygosity in single individuals ranged from 0.02 to 0.34 in the 24 accessions assigned to *M. acuminata* ssp. *banksii*. Inbreeding coefficient (F) confirmed significant excess of homozygous sites in these wild accessions, except for ‘AMB007’, ‘AMB008’ and ‘Sup04’ all collected in Maluku that were not originally classified as belonging to the *banksii* subspecies (Sutanto et al. 2016). It is noticeable that ‘AMB004’, also collected in Maluku has similar F and Het values than the accessions classified as ssp. *banksii*. In the ‘AA NG1’ cluster, heterozygosity ranged from 0.20 to 0.34 and inbreeding coefficient was reflecting excess of heterozygous sites for 22 accessions (**Table 2**).

**Table 2:** Heterozygosity (He) and inbreeding coefficient (F) for the accessions of the clusters ‘banksii’ and ‘AA NG1’

|  | ID | Name | N sites | F | Ho |
| --- | --- | --- | --- | --- | --- |
| **Cluster**<br>**'banksii'** | AMB004 | NA | 13538 | 0.84 | 0.02 |
|  | AMB007 | Utang/Biji | 15269 | -1.09 | 0.32 |
|  | AMB008 | Utang/Biji | 15247 | -0.96 | 0.30 |
|  | ITC0341 | Banksii | 12862 | 0.85 | 0.02 |
|  | ITC0428 | Higa | 9875 | 0.86 | 0.02 |
|  | ITC0453 | Banksii | 14587 | 0.59 | 0.06 |
|  | ITC0606 | Hybrid | 15136 | 0.83 | 0.03 |
|  | ITC0616 | Hawain 2 | 13231 | 0.86 | 0.02 |
|  | ITC0617 | Hawain 3 | 15262 | 0.84 | 0.02 |
|  | ITC0619 | Banksii | 14194 | 0.78 | 0.03 |
|  | ITC0620 | Banksii | 14896 | 0.87 | 0.02 |
|  | ITC0621 | Banksii | 14553 | 0.76 | 0.04 |
|  | ITC0623 | Banksii | 14480 | 0.86 | 0.02 |
|  | ITC0766 | Paliama | 14706 | 0.87 | 0.02 |
|  | ITC0806 | Banksii | 15156 | 0.88 | 0.02 |
|  | ITC0853 | Banksii | 14592 | 0.88 | 0.02 |
|  | ITC0867 | Banksii | 15236 | 0.78 | 0.03 |
|  | ITC0879 | Musa ac. ssp. banksii | 14872 | 0.87 | 0.02 |
|  | ITC0885 | Musa ac. ssp. banksii | 9800 | 0.78 | 0.03 |
|  | ITC0897 | Musa ac. ssp. banksii | 13061 | 0.23 | 0.12 |
|  | ITC0937 | Banksii | 13272 | 0.86 | 0.02 |
|  | ITC0955 | Banksii | 14695 | 0.87 | 0.02 |
|  | ITC1219 | Banksii | 14867 | 0.84 | 0.02 |
|  | Sup4 | Utang/Biji? | 13585 | -1.24 | 0.34 |
| **Cluster**<br>**'AA NG1'** | AROB003 | Mero Mero | 25904 | 0.13 | 0.20 |
|  | AROB004 | Wiau | 25010 | -0.15 | 0.27 |
|  | AROB005 | Duma | 24182 | 0.09 | 0.21 |
|  | AROB016 | Nape'e | 25744 | -0.03 | 0.24 |
|  | AROB019 | Tavilo | 26078 | -0.33 | 0.31 |
|  | AROB022 | Kararu 2 | 24989 | 0.06 | 0.22 |
|  | AROB034 | Nesuri | 26104 | -0.04 | 0.24 |
|  | AROB035 | Talasea | 26122 | -0.01 | 0.24 |
|  | AROB047 | Tobaung | 25856 | -0.16 | 0.27 |
|  | AROB049 | Nono 2 | 25954 | -0.27 | 0.30 |
|  | AROB050 | Sesévé | 25898 | -0.01 | 0.23 |
|  | ITC0589 | Gulum | 25181 | -0.44 | 0.34 |
|  | ITC0600 | Waimara | 16022 | 0.09 | 0.21 |
|  | ITC0770 | Navaradam | 23952 | -0.01 | 0.24 |
|  | ITC0773 | Mpiajhap | 19885 | 0.04 | 0.22 |
|  | ITC0778 | Gorop | 25711 | 0.08 | 0.21 |
|  | ITC0784 | Tamai | 25035 | -0.02 | 0.24 |
|  | ITC0798 | Garunga | 19648 | -0.20 | 0.28 |
|  | ITC0818 | Enar | 24672 | 0.02 | 0.23 |
|  | ITC0847 | Hova | 16980 | 0.05 | 0.22 |
|  | ITC0923 | Yapu Yapu | 24979 | -0.16 | 0.27 |
|  | ITC0929 | Loibwa | 19807 | 0.09 | 0.21 |
|  | ITC0949 | Wiliman | 25988 | -0.27 | 0.30 |
|  | ITC0984 | Yangun Yefan | 24766 | -0.02 | 0.24 |
|  | ITC1001 | Bogia Mun | 25866 | -0.03 | 0.24 |
|  | ITC1013 | Sena | 25660 | 0.09 | 0.21 |
|  | ITC1023 | Taoaya | 19889 | -0.23 | 0.29 |
|  | ITC1206 | Spiral | 26107 | -0.01 | 0.24 |
|  | ITC1220 | Yalumia | 25961 | -0.15 | 0.27 |
|  | ITC1244 | Mapua | 25848 | 0.10 | 0.21 |
|  | ITC1245 | Papat | 25459 | -0.48 | 0.34 |

For 24 accessions assigned to the subspecies *banksii*, we identified 1059 windows of 200kb size with more than 1 variable SNP. For each window, Tajima’s D values were plotted against nucleotide diversity (π) (**Fig. 4A**). Mean Tajima’s D in these windows was −0.28 (Variance = 1.28) and the distribution was skewed towards negative values, reflecting an excess of low frequency variants (**Fig. 4A, B and C**). The ‘AA NG1’ cluster has a greater diversity as expressed by nucleotide diversity (π) and the highest number of 200kb windows with more than 1 variable SNP (1502). For ‘AA NG1’, mean Tajima’s D parameter was 0.50 and the distribution of the values obtained for the windows was somewhat bimodal (variance = 2.47) with the main peak being largely negative, reflecting excess of low frequency variants, that could reflect purifying selection. The second peak is largely positive, reflecting excess of common variants that could be resulting from balancing selection (**Fig. 4B and C**). Windows with Tajima’s D below the 1% lower limit (−2.39) and above the 1% upper limit (3.72) for the cluster ‘AA NG1’ are presented in **Table 3**.

**Fig. 4:**
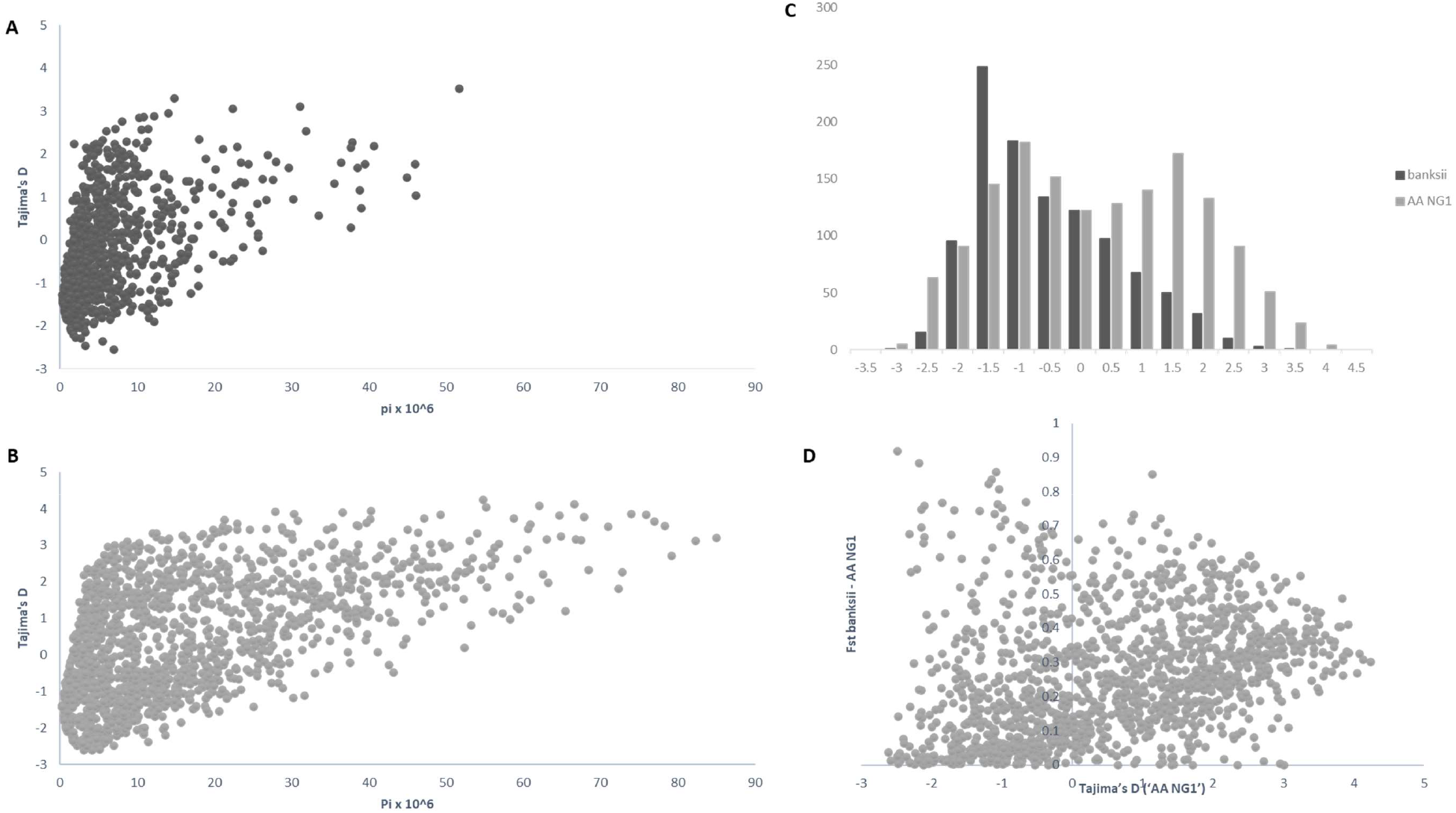
Distribution of pi (π), Tajima’s D and Fst calculated on 200 kb windows along the genomes for the clusters ‘banksii’ and ‘AA NG1’. Tajima’s D plotted against pi (π) for ‘banksii’ (A) and ‘AA NG1’ (B); comparative distribution of Tajima’s D values (C) and Tajima’s D calculated for ‘AA NG1’ plotted against Fst between ‘banksii’ and ‘AA NG1’ (D). Calculation were performed using VCFtools (Danecek et al. 2011).

**Table 3:** Windows (200 kb) with the 1% lowest and 1% highest values of Tajima’s D calculated for ‘AA NG1’, number of SNPs, Pi (π) and Fst between the clusters ‘banksii’ and ‘AA NG1’ are also displayed for these windows.

| Chrom. | Window | 'AA NG1' |  |  | 'banksii' |  |  | Fst |
| --- | --- | --- | --- | --- | --- | --- | --- | --- |
|  |  | SNPs | Pi | Taj.'s D | SNPs | Pi | Taj.'s D |  |
| chr01 | 11000000 | 40 | 73.95 | 3.85 | 27 | 12.25 | -1.78 | 0.31 |
| chr01 | 11200000 | 24 | 49.32 | 3.84 | 11 | 6.30 | -1.33 | 0.32 |
| chr01 | 11400000 | 15 | 30.28 | 3.85 | 17 | 9.08 | -1.30 | 0.26 |
| chr02 | 23800000 | 32 | 66.65 | 4.13 | 26 | 14.08 | -1.46 | 0.31 |
| chr02 | 24000000 | 26 | 54.80 | 4.24 | 29 | 11.57 | -1.81 | 0.30 |
| chr02 | 24200000 | 30 | 55.18 | 4.05 | 31 | 16.98 | -1.24 | 0.27 |
| chr02 | 24400000 | 25 | 47.13 | 3.73 | 15 | 6.53 | -1.81 | 0.36 |
| chr02 | 24600000 | 29 | 62.04 | 4.08 | 29 | 12.24 | -1.91 | 0.33 |
| chr02 | 25000000 | 18 | 36.61 | 3.90 | 1 | 0.37 | 0.00 | 0.33 |
| chr06 | 1800000 | 35 | 67.84 | 3.79 | 2 | 0.60 | -1.21 | 0.40 |
| chr10 | 21800000 | 33 | 64.73 | 3.81 | 3 | 6.15 | 1.65 | 0.40 |
| chr10 | 28400000 | 22 | 40.21 | 3.72 | 3 | 1.66 | -0.71 | 0.35 |
| chr10 | 28600000 | 20 | 40.28 | 3.93 | 3 | 1.88 | -0.50 | 0.33 |
| chr10 | 29600000 | 30 | 58.82 | 3.74 | 2 | 1.84 | -0.22 | 0.29 |
| chr11 | 1 | 40 | 75.93 | 3.84 | 2 | 2.30 | 0.15 | 0.49 |
| chr11 | 4800000 | 14 | 27.89 | 3.93 | 0 | 0.00 | 0.00 | 0.41 |
| chr01 | 6200000 | 33 | 6.95 | -2.48 | 5 | 3.29 | -0.75 | 0.14 |
| chr02 | 19800000 | 21 | 3.29 | -2.58 | 1 | 0.24 | 0.00 | -0.01 |
| chr03 | 29600000 | 29 | 5.84 | -2.40 | 2 | 1.14 | -0.61 | 0.08 |
| chr03 | 34400000 | 27 | 5.53 | -2.45 | 2 | 0.79 | -1.00 | 0.01 |
| chr04 | 3200000 | 15 | 2.25 | -2.48 | 2 | 1.67 | -0.38 | 0.02 |
| chr04 | 3400000 | 17 | 2.88 | -2.48 | 3 | 1.30 | -1.25 | 0.00 |
| chr04 | 5000000 | 22 | 4.20 | -2.48 | 3 | 4.92 | 1.42 | 0.92* |
| chr06 | 9800000 | 21 | 3.15 | -2.60 | 4 | 1.49 | -1.18 | 0.00 |
| chr06 | 10000000 | 26 | 4.94 | -2.46 | 3 | 3.50 | 0.62 | 0.03 |
| chr06 | 10200000 | 26 | 4.92 | -2.49 | 4 | 3.10 | -0.46 | 0.03 |
| chr06 | 10400000 | 29 | 5.19 | -2.59 | 4 | 2.93 | -0.74 | 0.01 |
| chr06 | 11600000 | 26 | 4.16 | -2.62 | 5 | 2.74 | -0.55 | 0.04 |
| chr06 | 17400000 | 13 | 1.93 | -2.42 | 0 | 0.00 | 0.00 | -0.01 |
| chr08 | 7200000 | 20 | 3.47 | -2.52 | 2 | 0.45 | -1.46 | -0.01 |
| chr08 | 38000000 | 45 | 11.40 | -2.39 | 4 | 3.65 | -0.32 | 0.06 |
| chr11 | 27800000 | 16 | 2.89 | -2.43 | 1 | 0.23 | 0.00 | -0.01 |

Whole genome Fst calculated between ssp. *banksii* and ‘AA NG1’ was 0.30. Considering 200 kb windows exhibiting more than 1 polymorphic SNP, the highest Fst value was 0.92 calculated on 25 SNPs on chromosome 4 (bin start 5.000.000). This genomic region also displayed the lowest Tajima’s D value and is likely under selection. However, plot of Tajima’s D values calculated for ‘AA NG1’ against Fst shows that most windows with negative Tajima’s D exhibits also low Fst (**Fig. 4D** and **Table 4**).

**Table 4:** Windows (200 kb) with 1% outliers for Fst calculated between the clusters ‘banksii’ and ‘AA NG1’. Number of SNPs, Tajima’s D and Pi are also displayed for these windows and for each cluster.

| Chrom<br>. | Window<br>(start) | SNP<br>s | Fst | ‘AA NG1’ |  |  | ‘banksii’ |  |  |
| --- | --- | --- | --- | --- | --- | --- | --- | --- | --- |
|  |  |  |  | SNPs | Taj.'s D | Pix10e6 | SNPs | Taj.'s D | Pix10e6 |
| chr01 | 12800000 | 6 | 0.77 | 0 | 0.00 | 0.00 | 6 | 1.26 | 8.11 |
| chr01 | 27400000 | 8 | 0.74 | 6 | -1.69 | 1.55 | 6 | 0.17 | 5.49 |
| chr04 | 4600000 | 10 | 0.77 | 10 | -1.85 | 3.22 | 1 | 0.00 | 0.44 |
| chr04 | 4800000 | 16 | 0.75 | 16 | -2.15 | 4.14 | 1 | 0.00 | 1.49 |
| *chr04* | *5000000* | 25 | *0.92* | 22 | *-2.48\** | *4.20* | 3 | *1.42* | *4.92* |
| chr04 | 5200000 | 15 | 0.88 | 14 | -2.18 | 3.45 | 3 | 0.25 | 2.62 |
| chr04 | 20400000 | 5 | 0.85 | 2 | 1.13 | 3.20 | 4 | -1.72 | 1.12 |
| chr04 | 20600000 | 3 | 0.88 | 1 | 0.00 | 1.86 | 3 | -1.52 | 0.87 |
| chr04 | 21200000 | 4 | 0.86 | 3 | -1.09 | 1.08 | 4 | -1.08 | 2.02 |
| chr04 | 21400000 | 5 | 0.76 | 5 | -1.06 | 2.66 | 5 | -0.90 | 3.41 |
| chr04 | 22000000 | 9 | 0.81 | 9 | -1.04 | 4.57 | 5 | -1.08 | 3.00 |
| chr04 | 30600000 | 22 | 0.76 | 19 | -2.10 | 5.90 | 21 | 0.93 | 26.70 |
| chr05 | 8400000 | 6 | 0.77 | 6 | -0.66 | 4.05 | 6 | -0.08 | 5.12 |
| chr06 | 13000000 | 17 | 0.75 | 16 | -1.00 | 10.54 | 9 | -1.35 | 4.41 |
| chr09 | 37800000 | 23 | 0.83 | 18 | -1.15 | 9.94 | 10 | 0.58 | 9.91 |
| chr09 | 38000000 | 34 | 0.82 | 24 | -1.19 | 13.09 | 21 | -0.31 | 16.60 |

## Discussion

### Within taxa diversity of *M. acuminata*

The analysis of the pruned set of 158 wild and cultivated accessions confirmed the assignation of the wild specimen to their respective species or subspecies in line with previous studies (e.g. Carreel et al. 1994; Perrier et al. 2011; Sardos, Perrier et al. 2016; Christelova et al. 2017; Martin, Cardi, et al. 2020). In addition, we detected introgressions of ssp. *burmannica/siamea* into several *M. acuminata* ssp. *malaccensis* samples from Thailand, confirming genetic contacts between both genepools in the region (Rouard et al. 2018; Martin, Cardi, et al. 2020). Interestingly, we revealed similar genomic compositions of ‘Borneo’ classified as ssp. *microcarpa* and the two ssp. *errans* from the Philippines. Martin, Cardi, et al. (2020) considered the ‘Borneo’ accession as a significant contributor to a ‘banksii’ genepool without any traces of admixture. Our wider sample rather suggests a partially common genetic background between ssp. errans/’Borneo’ and ssp. *banksii* but with additional variability specific to ssp. errans/’Borneo’. We considered in our study ssp. *banksii* and ssp. errans/’Borneo’ as two separated genepools.

Resolving the origins and the evolution of *Musa acuminata* genepools, in particular of those linked to domestication, has been an active area of research. Studying sequences of four genic regions in a wide sample of Musaceae, Janssens et al. (2016) inferred *Musa* section (including *M. acuminata* and *M. schizocarpa*) dispersal history. According to these findings, the origins of *Musa* section bananas are in the northern Indo-Burma region. First migration occurred towards what is today the Malayan peninsula and Sumatra via land. From there, migrations occurred to Java, Borneo, the Philippines, New-Guinea island and, lastly, back to the southern Indo-Malayan region. If this model applies broadly to the *Musa* species belonging to the section *Musa*, the study of Rouard et al. (2018) using whole genome sequence of four *M. acuminata* subspecies, ssp. *burmannica, zebrina, banksii* and *malaccensis* to infer phylogeny at the intra-species level suggested similar radiation pattern. The distribution of the diversity of the *M. acuminata* subspecies as inferred in the present study is also congruent with this dispersion scenario (**Fig. 2**). The role of the Malayan peninsula and Sumatra as a centre of secondary diversification and radiation (Janssens et al. 2016) could also explain the highly admixed profiles of the subspecies *sumatrana* and *truncata*, located north-west of Sumatra and in the Malayan peninsula mountains respectively. In the absence, in the sample, of populations for these taxa, these high degrees of admixture inferred by the software rather reflect shared ancestry with different genepools than introgressed profiles.

### Undefined ancestral genepools in south east Asia

The Bayesian Markov Chain Monte Carlo (MCMC) approach was demonstrated efficient in identifying source populations for which unadmixed individuals are absent in the sample (François and Durand 2010). This is confirmed in our study where two genepools that appeared specific to cultivated bananas, ‘Und-I’ and ‘Und-III’, were both present into ‘Pisang Madu’. This is in accordance with Martin, Cardi, et al. (2020) who identified two cryptic ancestor populations co-existing within an accession also called ‘Pisang Madu’ (of same origin). In addition, the analysis of our wider sample suggests that one of this genepool, ‘Und-I’, is widely present within cultivated diploid bananas, especially those originating in SEA and confirmed previous studies (Carreel et al. 2002; Jeensae et al. 2021). The presence of three cultivated accessions from Thailand inferred with more than 90% of their genome belonging to this cluster suggests a Thai origin to ‘Und-I’ (**Fig. 3**). The genepool ‘Und-III’ identified in ‘Pisang Madu’ and ‘Pisang Jari Buaya’ from the Indo/Malaysian region may correspond to the rare mitotype β identified by Carreel et al. (2002). The collection sites of these accessions plead for a potential origin of ‘Und-III’ in island SEA rather than on the continent, but further investigation should be performed (**Fig. 3**).

In congruence with Perrier et al. (2011), the genomic constitutions inferred for edible diploids showed high levels of admixture that follow the routes of linguistic diffusion in both directions. At the extremes of the species range, the eastern ssp. *banksii* signature was identified in the sole AA from India and as far as in East Africa while the Myanmar ssp. *burmannica/siamea* at the west was detected as introgressions in accessions from the Philippines but was not identified in any of the AAs from NG (**Fig. 3**).

However, despite the predominantly admixed profiles of edible AAs, the analysis presented suggested the occurrence of accessions in SEA with no introgression from ssp. *banksii* and a group of accessions in NG that may be not introgressed by any of the *M. acuminata* subspecies from SEA. It was consistent with findings obtained with DArT and GBS-derived SNPs markers (Sardos, Perrier et al. 2016; Sardos, Rouard et al. 2016). Given that all three analyses, including the present one, were performed on reduced sets of markers that may have failed detecting small introgressions, such as the one inferred by Martin, Cardi, et al. (2020) in some of the cultivated accessions. We therefore used the Patterson’s D test to further confirm or infirm the absence of introgressions in these accessions (**Supplementary Tables S2** and **S3**, supplementary material Online). The absence in the sample of the taxon of origin (P1), likely corresponding to the ‘Und-I’ genepool, for most of the cultivated accessions from SEA, hampered the test to be conclusive. Equally, and even though some of the results obtained for a few PNG accessions suggested no introgressions by the Asian wild taxa tested, incoherence was observed in the BBAA, ABBA and BABA patterns counts that questions the choice of the outgroup, *M. schizocarpa*, at least for some of the accessions tested.

### The origin of bananas in New Guinea island

The prevalence in the sample of accessions originating in NG island both for the wild specimen and the cultivated diploids constitute a unique opportunity to compare wild and cultivated populations evolving in the same environment. We therefore compared the closely related *M. acuminata* ssp. *banksii* and the cluster ‘AA NG1’. In domesticated crops, selection is expected to induce either an excess of low frequency polymorphism due to post-domestication’s bottleneck expansion, as in selfing chickpea (Varshney et al. 2019), or a drop in rare alleles frequencies due to recent selection, such as in clonal African yams (Akakpo et al. 2017). However, the signal is not as clear in the cluster ‘AA NG1’ as it suggests both types of phenomenon with potentially both positive selection and balancing selection at work, such as in vine, another clonal crop (Houel et al. 2010). More importantly, despite a common genetic background, the AAcv from the cluster ‘AA NG1’ revealed higher levels of diversity than the wild *M. acuminata* ssp. *banksii* (**Fig. 4**). It differs substantially from standard wild-to-domesticate transition scenario in which domestication is expected to induce a loss of genetic diversity (Meyer and Purugganan 2013). Interestingly, same pattern was observed in sugarcane, another clonal crop from NG, in which two mechanisms were highlighted, balancing selection and the accumulation of mutations due to clonal propagation (Arro et al. 2016). Both could apply to the ‘AA NG1’ cluster. However, undetectable introgressions of one or more genepools not present in the sample cannot be ruled out. In fact, the existence of an unexplored genepool of *M. acuminata* in Indonesian New Guinea is suspected for a long time (Simmonds 1956; Argent 1976). In such context, the *M. acuminata* collected in the Wallacean islands of Ambon and Seram, at the west of NG, are especially interesting as three of them exhibited slightly different genetic backgrounds coupled to higher levels of heterozygosity than the ssp. *banksii* (**Fig. 2** and **Table 2**).

On the other hand, *M. acuminata* ssp. *banksii* in NG showed a clear signature of a population under expansion after a bottleneck. It is compatible with the hermaphrodism of many of the female flowers observed in this subspecies, a feature that is not seen in other *M. acuminata* subspecies at the exception of ssp. *errans* in the Philippines and a form of ssp. *burmannica* in India (Simmonds 1956; Argent 1976; Joe et al. 2016). Bottleneck can be due to the establishment of selfing (Foxe et al. 2009; Guo et al. 2009) but it could also jointly be resulting from the last glacial period during which the climate was cooler and drier (Bowler et al. 1976; Hope et al. 2004), inducing conditions less favourable for *M. acuminata* in NG. The northern coast lowlands were likely the only rainforest refugee of the island at the time (Evans 2020). Stabilization in climate, estimated 9,000 years ago, interestingly roughly coincided with the moment when humans of the region were on their way to domesticate local crops, with clear evidence of established agricultural activities in NG highlands dated back at 7,000 years ago (Denham et al. 2003). However, the switch from gathering to agriculture was not a discrete event but a gradual process that was preceded by a range of intermediate practices (Denham et al. 2003; Summerhayes et al. 2017). Some forms of management of natural populations may have started on coastal territories, more suitable for human settlements during the last glacial period. As climate was warming and sea level was rising, these practices may have been gradually imported inland, along with exploited plants, as humans were leaving the coasts. Alternatively, a mountain form of *M. acuminata*, nowadays extinct or undiscovered, may have survived in one of the separated glacial refugees located in the many mountains’ valleys of the island. These refugees are believed to be responsible for the high floral variability of NG mountain forests (Hope et al. 2004). This mountainous *M. acuminata* could have been exploited by humans who rapidly settled in the mountains when conditions became more favourable (Summerhayes et al. 2017). This second scenario could explain the presence of *Musa* section seeded bananas in pre-cultivation archaeological records in NG highlands, an unsuitable environment for present days wild *Musa* section bananas (Denham et al. 2003).

Under both perspectives, it seems reasonable to envisage that present day ssp. *banksii* may have been highly influenced by past human populations. Especially since archaeological evidence of past cultivation of seeded forms of *Musa* bananas were discovered in the highlands of NG (Denham et al. 2003), where *ssp. banksii* does not grow. Similar archaeological remains were identified in the Bismarck archipelago and Vanuatu that are both outside the natural range of the species, pleading for a human introduction (Lentfer 2009; Tromp et al. 2020). The existence of *M. acuminata* ssp. *banksii* exhibiting some levels of parthenocarpy, an essential trait for nowadays cultivated bananas, in the far Samoa (Simmonds 1956; Sardos, Sachter-Smith, et al. 2019) may result from the same dispersal wave. Since these Samoan plants, now growing feral, fall genetically within ssp. *banksii* from NG when genotyped with SSR (Hribova pers. com.), it may well be that what we know nowadays as *M. acuminata* ssp. *banksii* has undergone some form of human selection in the past. In such case, whether the source population of present days spp. *banksii* still exist is unresolved but its possible existence echoes the suspected presence of a population with a wider genetic diversity in some unexplored region of NG island.

## Conclusion

By studying the edible diploid bananas with an AA genomic composition, we aimed at deciphering the first stage of banana domestication, namely the transition from *M. acuminata* to edible AAs. Doing so, we confirmed the existence in edible AAs of two genepools for which no corresponding wild taxa was identified. The results suggest Thailand as a candidate region of origin for one of these genepools. The low number of accessions introgressed with the second undefined genepool may suggest contribution from germplasm outside *M. acuminata* species. Therefore, if such gene pool has subsisted, wild species growing in a region from the Malaysian peninsula, Borneo and the Philippines should be investigated. In Papua New Guinea, *M. acuminata* ssp. *banksii*, an important contributor to edible bananas, showed clear evidence of a population that underwent a strong bottleneck. It could be due jointly to last glacial period and to the establishment of selfing in the taxa. Humans, who impacted flora and fauna in the region (Fairbairn et al. 2006), could be an additional factor. In this context the higher levels of diversity discovered in the edible AA closely related to this subspecies are surprising. Especially since no apparent traces of introgressions were identified in many of these cultivated samples. We therefore questioned the existence of an undiscovered source population for both *M. acuminata* ssp. *banksii* and closely related edible AA in NG.

## Materials and methods

### Plant materials

A set of 226 diploid banana accessions was selected (**Table 1**; Supplementary Table S1). This set comprised 68 wild accessions belonging to subspecies of *M. acuminata*, 154 related edible diploid cultivars, three accessions of *M. schizocarpa* considered as outgroup and a hybrid between *M. acuminata* ssp. *banksii* and *M. schizocarpa*. These materials were provided by the ITC (170 samples Musa Germplasm Information System (MGIS - https://www.crop-diversity.org/mgis/) (Ruas et al, 2017), the banana collecting mission to the AROB (25 samples) (Sardos et al. 2018), CIRAD (24 samples) (Perrier et al. 2019), collecting missions to Indonesia (4 samples) (Sutanto et al. 2016) and EMBRAPA (3 samples).

### Restriction-site-associated DNA sequencing

DNA from each accession was extracted following a 2X CTAB protocol (modified from Doyle and Doyle 1990). Library for restriction-site-associated DNA sequencing (RADSeq) was built with the PstI restriction enzyme. The 300–500 short-insert libraries were sequenced with 91 bp paired-end reads using Illumina HiSeq2000 (Illumina, San Diego, CA, USA) by BGI Hong Kong. At BGI, the raw data were modified with the following two steps: (1) reads polluted by adapter sequences were deleted; and (2) reads that contained >50 % low-quality bases (quality value ≤5) or >10 % N bases were removed.

### Read processing and SNP calling

Reads contained in raw FASTQ files (one per sample) were checked using FastQC and then cleaned to remove Illumina adapter sequences and low-quality ends (Phred score > 20) with Cutadapt (Martin 2011). After trimming, reads inferior to 30 bp were discarded. MarkDuplicates from Picard tools (Picard Toolkit 2019) was used to remove tags duplicate reads. Reads were then aligned against the *Musa acuminata* genome v2 downloaded on the Banana Genome Hub (Droc et al. 2013) using BWA-MEM (Li and Durbin 2010). Re-alignment was done with the IndelRealigner module from GATK v4.1 (McKenna et al. 2010). We then followed the GATK pipeline recommended for a non-model organism by adding the recalibration step. It consisted in performing an initial round of SNP calling on the original uncalibrated data, selecting the SNPs with the highest confidence, and then executing a round of base recalibration on the original mapped reads files. The SNP calling was done with the GATK module HaplotypeCaller v4.1 to call SNPs and indels. Considering inter-sample variation, the SNP calling was done on all samples simultaneously. The pipeline used to perform those analyses is available at https://github.com/CathyBreton/Genomic_Evolution.

### Genetic diversity analyses

For the initial set of 226 accessions, SNPs were filtered for missing data (5% as maximum allowed) and MAF (1%), yielding a total of 39,031 bi-allelic SNPs. At this stage, eight accessions were discarded as presenting more than 15% of missing data.

A dissimilarity matrix was calculated following the Simple-Matching index with a minimum of 70% of common sites between each pair of remaining individuals with DARwin 6 (Perrier and Jacquemoud-Collet 2006). A weighted Neighbor-Joining (NJ) tree rooted using *M. schizocarpa* as outgroup was then constructed. This first tree allowed identifying cultivated accessions exhibiting identical or nearly identical genotypes and corresponding to duplicates or clonal varieties. To avoid potential bias in further analyses due to genotypes redundancy, we then allowed the presence of a single accession per Genotype Clusters, further reducing the set of accessions to 158 individuals. For each Genotype Clusters, selection of the unique representative kept for further analyses was based on the lowest rate of missing data per accession. For this subset of 158 representative accessions, we retrieved a new set of SNPs with a minor allele frequency of 0.01 (1%) or greater and allowing a maximum of 10% of missing data. With the 66,481 SNPs obtained, DARwin 6 was used to calculate a simple-matching distance matrix. A new weighted neighbour-joining tree rooted on *M. schizocarpa* was then constructed on the set of 158 accessions, eliminating potential distortion due to duplicated accessions.

### Global population structure

Using VCFtools (Danecek et al. 2011), we generated a set of SNPs evenly distributed every 100 kb and not allowing missing data to reflect all chromosomal regions. This set involved 1278 SNPs used to investigate the structure of the 158 accessions of the pruned dataset. We used a Bayesian Markov Chain Monte Carlo (MCMC) approach implemented in the program STRUCTURE v2.3 (Pritchard et al. 2000). The admixture model with the assumption of correlated allele frequencies between groups (Falush et al. 2003) was chosen and 5 replicates of each value of k ranging from 1 to 15 were run with a burn-in-length of 50,000 followed by 150,000 iterations of each chain. The most likely true of the values of k was determined by examining DeltaK, an *ad hoc* quantity related to the second order rate of change of the log probability of data with respect to the number of clusters (Evanno et al. 2005) and plotted using STRUCTURE HARVESTER (Earl and vonHoldt 2012). STRUCTURE was then run again following the same model for the best values of K identified with 5 replicates each and a burn-in length of 200,000 followed by 800,000 iterations of each chain.

### Tests for introgression in cultivated bananas (AA)

The four taxon Patterson’s D test (Green et al. 2010; Durand et al. 2011) was developed to detect introgressions in closely related taxa. It considers an ancestral “A” allele and a derived “B” allele across the genome of four taxa with a tree topology (((T1,T2),T3),O). Under the hypothesis “without introgression” the two allelic patterns at the tip of the tree, “ABBA” or “BABA”, occur with equal frequency. An excess of “ABBA” or “BABA”, reflected by a D-statistic significantly different from zero, indicates potential gene flow between P2 and P3 or P1 and P3, respectively. Here, we used the derived statistic *f_d_* that is a more conservative estimator of introgression developed for small number of SNPs (Martin et al. 2015) implemented in https://github.com/simonhmartin/genomics_general. A R script allowed the calculation of the P-value based on jackknife for the null hypothesis that *f_d_* is 0. The procedure is available at https://github.com/CathyBreton/Genomic_Introgression_ABBA_BBAA_Test. Two tests were performed. In the first one, 31 Papuan edible AAs closely related to the Papuan wild *M. acuminata* ssp. *banksii* were tested for introgression by the subspecies originating in SEA. In the second one, 23 edible AA from SEA were tested for introgression by the Papuan *M. acuminata* ssp. *banksii*. Accessions selected as representative for each wild taxon are presented in **Supplementary Tables S2** and **S3** (Supplementary materials online).

### Population differentiation between wild and cultivated Papuan bananas

Based on STRUCTURE outputs, we considered a sub-cluster of 31 AA from PNG with a *M. acuminata* ssp. *banksii* genomic background over 90%. This population and its wild counterpart are represented in our samples by 31 and 24 accessions respectively. A set of 238,357 SNPs was retrieved from GIGWA (Sempéré et al. 2019; Rouard et al. in press) allowing a maximum of 50% of missing data for each of the two populations. Using VCFtools, we first assessed observed heterozygosity (Ho) and inbreeding coefficient (F) for each of these accessions. Then, considering 200 kb windows exhibiting more than one SNPs, we calculated the nucleotide diversity (π) and Tajima’s D for each population. Finally, we calculated weighted Fst between the cultivated accessions and their wild relative, genome-wide and for 200 kb windows along chromosomes. To better understand the nature of selection in AA NG1 we then considered the 1% lowest and highest Tajima’s D values and the 1% greater Fst values.

## Data availability

Raw sequence reads were deposited in the Sequence Read Archive (SRA) of the National Center for Biotechnology Information (NCBI) (BioProject: PRJNA450532). SNP datasets are available on GIGWA (Sempéré et al. 2019) via MGIS (https://www.crop-diversity.org/mgis/gigwa) (Ruas et al. 2017).

## Acknowledgements

We thank BGI for their technical assistance and services for the RAD sequencing. This work was technically supported by the CIRAD - UMR AGAP HPC Data Center of the South Green Bioinformatics platform (https://www.southgreen.fr/). This work was financially supported by CGIAR Fund, and in particular by the CGIAR Research Program, Roots, Tubers and Bananas and the Genebank platform.

