## Supplementary figures and images for "Wild to domesticates: genomes of edible diploid bananas hold traces of several undefined genepools"

### Supplementary Fig. S1

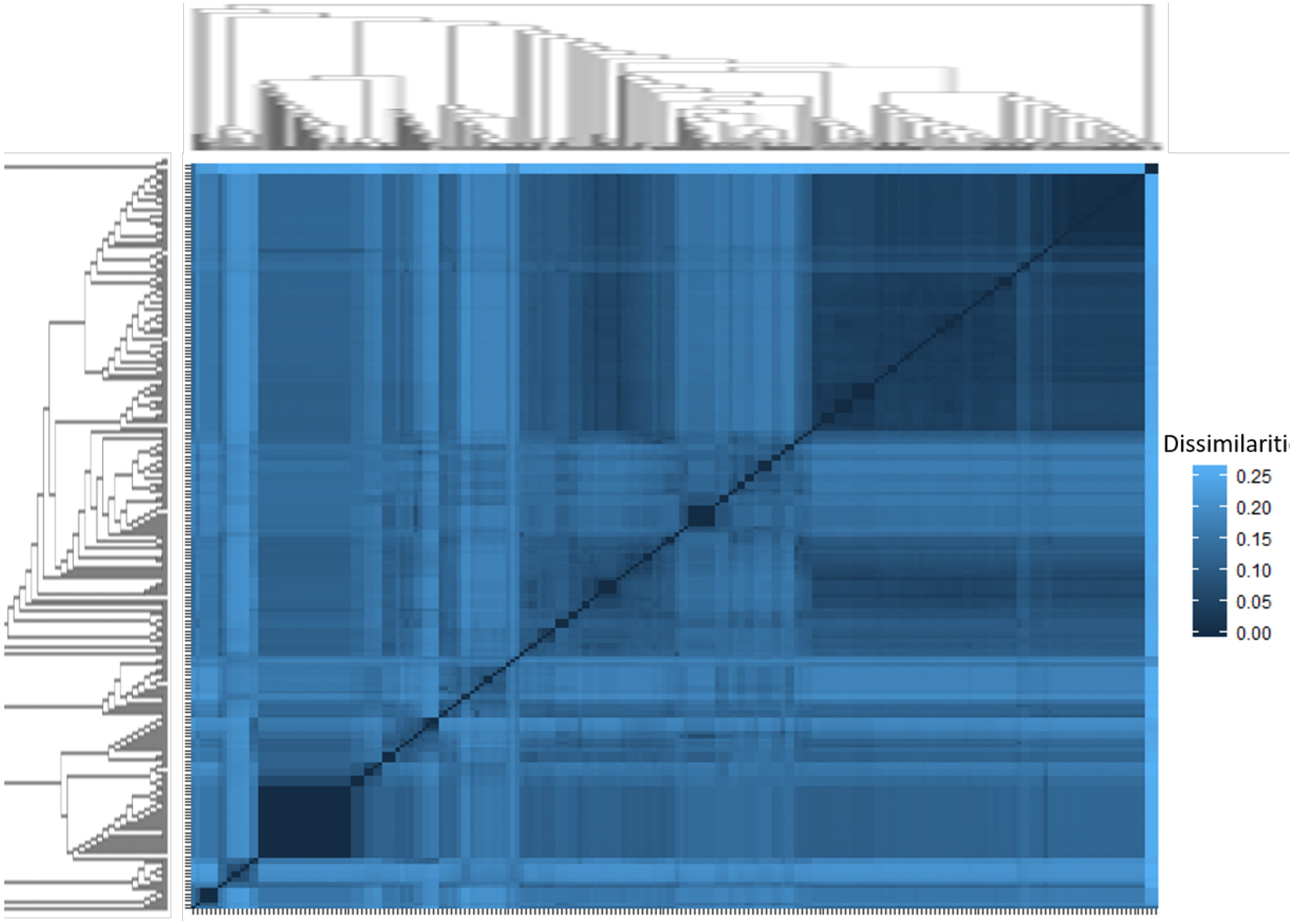
